# Reservoir ecosystems support large pools of fish biomass

**DOI:** 10.1101/2023.09.04.556263

**Authors:** Christine A. Parisek, Francine A. De Castro, Jordan D. Colby, George R. Leidy, Steve Sadro, Andrew L. Rypel

## Abstract

Humans increasingly dominate Earth’s natural freshwater ecosystems, and many freshwater fisheries resources are imperiled and at-risk of collapse. Yet despite this, the productive capacity of intensively modified freshwater ecosystems is rarely studied. We digitized, and provide open access to, a legacy database containing empirical fish biomass from 1,127 surveys on 301 USA reservoirs. In parallel, we developed a slate of reservoir classification schemas that were deployed to better understand distributions of biomass and secondary production. By fusing these data products, we generated a predictive capacity for understanding the scope of fisheries biomass and secondary production across all USA reservoirs. We estimate total potential fish total standing stock in USA reservoirs is 3.4 billion (B) kg, and annual secondary production is 4.5 B kg y^-1^. In southern USA alone, total standing stock and secondary production are 1.9 B kg and 2.5 B kg y^-1^, respectively. We also observe non-linear trends in reservoir fish production and biomass over time, indicating that these ecosystems are quite dynamic. Results demonstrate that reservoirs represent globally relevant pools of freshwater fisheries, in part due to their immense spatial and limnetic footprint. This study further shows that reservoir ecosystems play major roles in food security, and fisheries conservation, even though they are frequently overlooked by freshwater scientists. We encourage additional effort be expended to effectively manage reservoir environments for the good of humanity, biodiversity, and fishery conservation.

**Significance Statement:** Globally, many freshwater fishes and fisheries resources are imperiled and at-risk of collapse. However, previous research overwhelmingly focuses on freshwater fisheries in natural rivers and lakes. This study provides evidence that novel and reconciled ecosystems, such as reservoirs, hold massive pools of freshwater fisheries biomass and may have higher ecological value than previously thought. While dams are patently ecological catastrophes, ecosystem services including secondary fish production provided by reservoirs are nonetheless substantial. Indeed, in many locations (e.g., arid regions), reservoirs are the only remaining fisheries resource. We suggest considerable conservation management is warranted for reservoir fisheries worldwide.

**Data Deposition:** Code is available on GitHub (https://github.com/caparisek/res_biomass_USA; DOI 10.5281/zenodo.8316696). All data and reservoir classifications are available on Zenodo (DOI 10.5281/zenodo.8317007). Furthermore, data will also be deposited in the Environmental Data Initiative repository upon acceptance of this manuscript.

## Introduction

Human dominance over freshwater ecosystems highlights the necessity to understand the fragility and productive capacity of these natural resources in response to global environmental change (1, 2). Inland fisheries are especially critical, providing protein to developing countries (3, 4), cultural value (5, 6), and economic development (7, 8). Freshwater fisheries and diversity are under threat from a range of sources including overfishing, pollution, habitat fragmentation, invasive species, and climate change (9, 10). These alterations have prompted widespread declines in freshwater fisheries; trends unlikely to abate given current socioecological trajectories (11, 12). Furthermore, harvest of marine fisheries stocks has plateaued since 1989 (13, 14), suggesting additional fisheries resources will be needed to sustain human societies in the future. And while aquaculture is increasingly filling gaps, cultured fish are not currently scaling sustainably (15, 16). Inland fisheries will continue to be a major food source globally, but many inland fisheries are data-limited, presenting a challenge to conservation (17, 18).

Reservoirs represent potentially overlooked pools of secondary production (hereafter, “production”). Indeed, impoundments of streams and rivers by dams, are increasingly prominent features over landscapes (19, 20). Dams have altered over half of Earth’s large rivers, including eight of the most speciose ecosystems (19). Overall, dams decimate native fish diversity and other freshwater riverine communities (21–23). In a sobering assessment, Benke 1990 (24) estimated that only 42 high quality free flowing rivers remain in the contiguous USA. Species that persist in reservoirs tend towards remarkably similar faunas composed of resilient species, often characteristic of warm-water lakes (25). Yet production varies widely across reservoir ecosystems that differ in shape, residence time, temperature, depth, and other factors (26).

Further, many reservoirs may have declined in production as dams and reservoirs have aged towards or beyond expected lifespans (27–29). Reservoirs have historically been understudied, and little research exists on the distribution, limnology, and ecology of reservoirs (but see 30, 31), as instead many scientists opt for the study of more natural lakes. Yet the sheer surface area of impounded waters on Earth implies that these environments produce ecosystem services that we should study and manage for improved sustainability. For example, understanding the ecological value of reservoirs may be critical for adapting to future climate change and food security challenges.

The primary goals of this research are to: (**1**) Digitize and make publicly available a legacy database containing fish biomass estimates from USA reservoirs; (**2**) Develop a reservoir classification system with broad application for 85,470 USA reservoirs such that any reservoir can be placed within families of similar reservoir types; (**3**) Test the degree to which biomass in individual reservoirs and reservoir types has changed over long time periods; and (**4**) Generate fish biomass predictions in all USA reservoirs to a standardized point in time and estimate total biomass and annual production rates for fish populations across all USA reservoirs.

## Results

Fish biomass and production rates in 301 sampled USA reservoirs were highly variable in space and time. Across all **sampled reservoirs**, total standing stock predicted for the standardized year (1993) ranged 802 kg – 103 million (M) kg with mean standing stock for an average reservoir = 3.14 M kg (+/- 0.47 SE) (**Dataset S1**). Similarly, production rates across sampled reservoirs ranged 1,043 kg y^-1^ – 135 M kg y^-1^ with mean production for an average reservoir = 4.1 M kg y^-1^ (+/- 0.61 SE).

We created a series of reservoir classification schemas of ascending complexity that placed 85,470 USA reservoirs into families of reservoirs with similar underlying characteristics. Our most complex classification system (Schema 5) was highly detailed and combined data on ecoregion, total reservoir storage capacity (m^3^), and water discharge (m^3^s) from dams (**Dataset S2**). Out of five Generalized Additive Mixed Models (GAMMs), each of which applied one of the unique classification schemas, Schema 5 yielded the best model for making total standing stock predictions (**Figure 1; Figure S1-2; Table S1**). Thus, within any given ecoregion, four different clusters of reservoirs emerged: 1) small volume and low discharge; 2) small volume and high discharge; 3) large volume and low discharge; and 4) large volume and high discharge (**Figure 2**).

**Figure 1.**
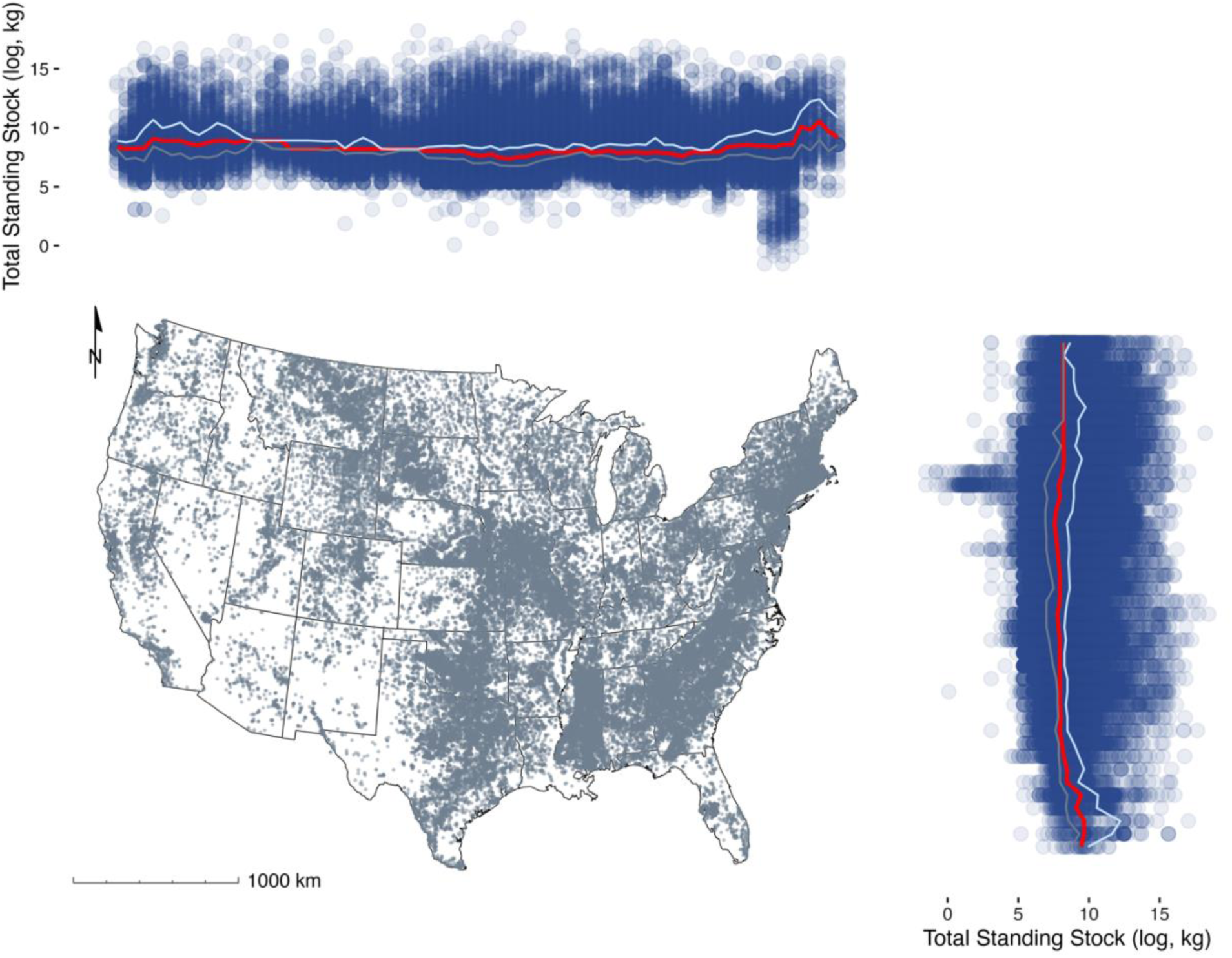
Map of contiguous United States representing all NID reservoirs (gray) and estimated total standing stock (log, kg) of those reservoirs by binned longitude and latitude (blue). Median (red), 25% quartile (gray), and 75% quartile (cyan) estimates by binned longitude and latitude total standing stock (log, kg) are overlaid.

**Figure 2.**
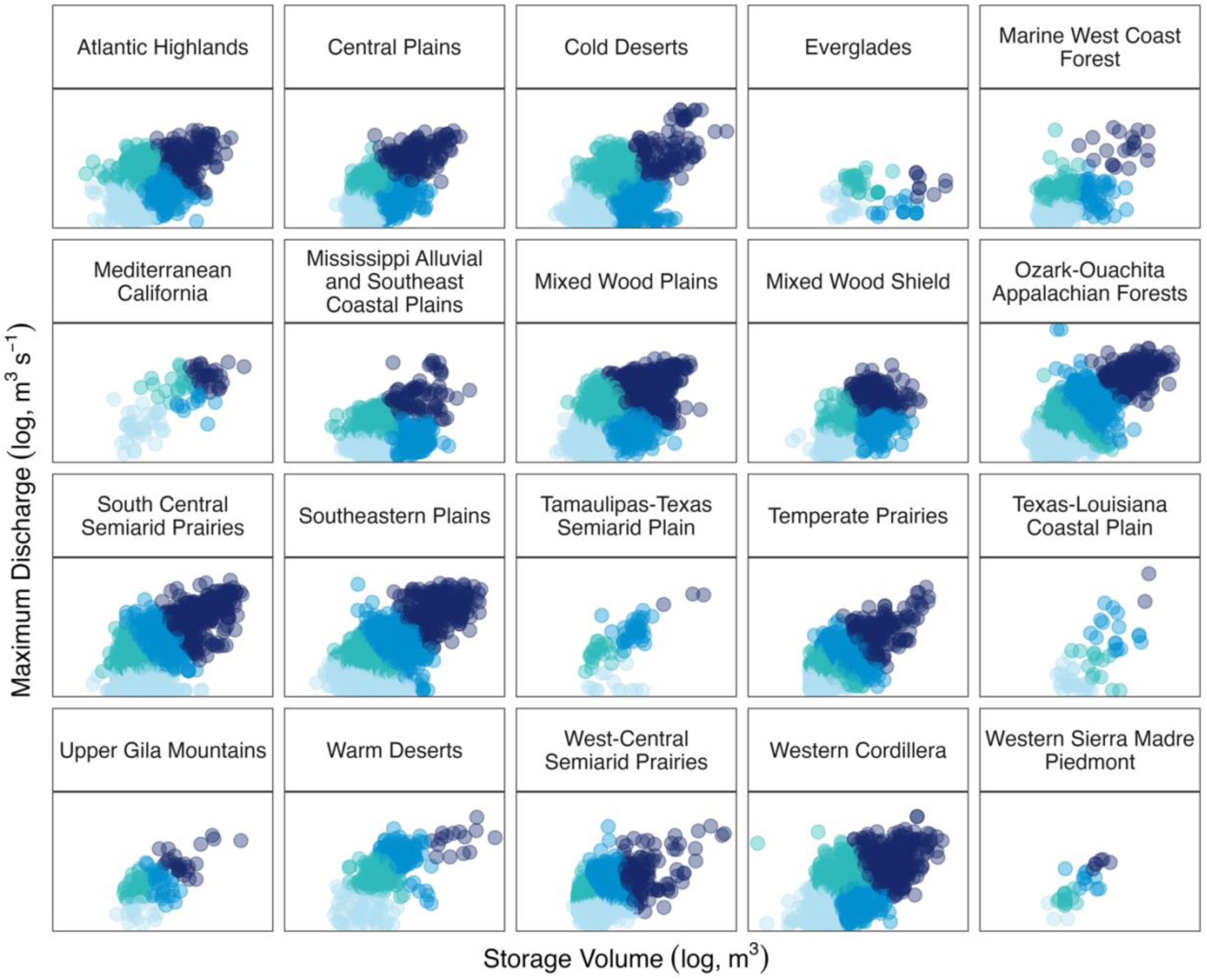
Results from k-means analysis by ecoregion using 4-cluster separation on USA reservoir discharge (log, m^3^ s^-1^) and storage volume (log, m^3^) (n = 36,340). Clusters represent reservoirs with small volume and low discharge (light blue), small volume and high discharge (turquoise), large volume and low discharge (medium blue), and large volume and high discharge (dark blue).

By combining empirical biomass data with our highest ranked reservoir classification system, we estimate **southern USA reservoirs (n = 55,731)** contain 1.92 billion (B) kg (+/- 0.09 SE across calculations) of fish mass, and total annual production for the region ranges 2.20 – 2.78 B kg y^-1^ (+/- 0.12 SE across calculations). Expanding to the **entirety of the USA (n = 85,470)**, we estimate total reservoir standing stock is 3.43 B kg (+/- 0.18 SE across calculations) with production ranging 3.87 – 5.01 B kg y^-1^ (+/- 0.23 SE across calculations) (**Table 1; Table S2; Dataset S3**). The top 5 USA states in total standing stock of reservoir fishes included Texas (368 M kg), Arkansas (187 M kg), Oklahoma (184 M kg), Florida (147 M kg), and South Dakota (147 M kg). Most states have reservoir standing stocks <100 M kg (**Figure 3A**); however when states are scaled by surface area, divergent state ranking patterns emerge. For example, Louisiana (611 kg ha^-1^), Indiana (607 kg ha^-1^), Alabama (601 kg ha^-1^), Maryland (592 kg ha^-1^), and Illinois (571 kg ha^-1^) had the highest mean biomasses, but none of these states were in the top five for total standing stock (**Figure 3B**). By Omernik **level II ecoregion**, we predicted total standing stock is highest in the following ecoregions: Southeastern Plains (950 M kg), Ozark-Ouachita Appalachian Forests (400 M kg), and South Central Semiarid Prairies (378 M kg), and lowest in: Western Sierra Madre Piedmont (0.93 M kg), Upper Gila Mountains (11 M kg), and Texas-Louisiana Coastal Plain (25 M kg). Incorporating reservoir storage and discharge, total predicted standing stock is greatest in large volume and high discharge reservoirs of: Southeastern Plains (695 M kg), Ozark-Ouachita Appalachian Forest (341 M kg), South Central Semiarid Prairie (303 M kg), and lowest in reservoirs with small volume from: Western Sierra Madre Piedmont reservoirs with low discharge (0.003 M kg) and high discharge (0.050 M kg), Tamaulipas-Texas Semiarid Plains reservoirs with high discharge (0.073 M kg) and low discharge (0.151 M kg), and Upper Gila Mountains reservoirs with high discharge (0.168 M kg).

**Figure 3.**
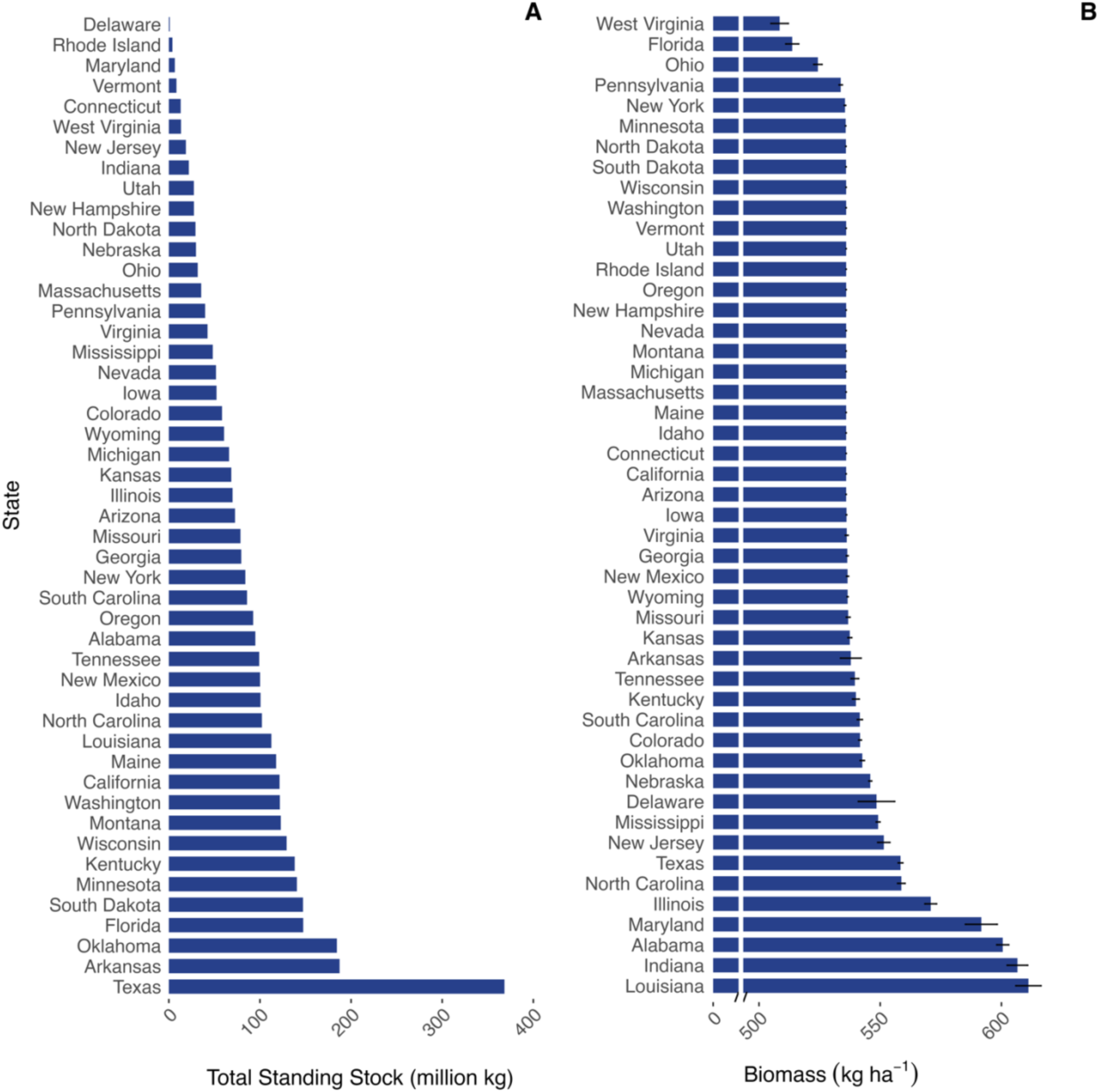
(**A**) Summary of total standing stock (million kg) estimates by state for reservoirs of the contiguous USA. (**B**) Mean fish biomass (kg ha^-1^) for each state which relates total standing stock (Panel A) to relative surface area of water available (ha) in that state; error bars represent the standard error of the mean.

**Table 1.** Total southern and all USA reservoir fish total standing stock and production estimates.

| Southern USA |  |  | USA |  |
| --- | --- | --- | --- | --- |
| Schema | Total Standing Stock (kg) | Production (kg y<sup>-1</sup>) | Total Standing Stock (kg) | Production (kg y<sup>-1</sup>) |
| – 1 –<br>Simple Average | 1,693,346,335 | 2,201,350,236 | 3,031,693,257 | 3,941,201,234 |
| – 2 –<br>Large & Small | 2,001,994,842 | 2,602,593,295 | 3,587,800,617 | 4,664,140,802 |
| – 3 –<br>Size-Flow | 1,713,033,098 | 2,226,943,027 | 2,976,459,235 | 3,869,397,006 |
| – 4 –<br>Ecoregion | 2,055,598,037 | 2,672,277,448 | 3,689,453,361 | 4,796,289,369 |
| – 5 –<br>Eco-Size-Flow | 2,137,464,393 | 2,778,703,711 | 3,855,651,138 | 5,012,346,479 |
| Mean | 1,920,287,341 | 2,496,373,543 | 3,428,211,522 | 4,456,674,978 |

Reservoir biomass and production are highly variable over long time periods. For example, we observe patterns of relatively constant biomass, increasing and declining biomass, and spikes in production followed by decreases that ultimately return to a baseline. Importantly, documentation of lake-specific trends allowed for standardization of biomass estimation to a given year of interest. Although we triangulated on one year (1993) for this analysis, this same technique could be applied to standardize biomass estimates to any year of interest, while still accounting for important lake-specific trends.

We collated additional data on forty-two independent poisoning surveys for USA reservoirs that were not part of the National Reservoir Research Program legacy dataset as a validation dataset (32, 33). A mixed effect regression model using classification method as a random effect showed that total standing stock from independent surveys was strongly correlated with predicted total standing stock values from the same reservoir (**Figure S3**). Furthermore, the slope of this model = 0.98 and R^2^c (pseudo-R^2^ for both fixed and random effects) = 0.98, expressing a near one-to-one relationship that did not differ significantly from a slope = 1. Further validation showed trends between observed fish biomass as a function of predicted fish biomass also followed a line with slope = 1 and intercept = 0 (**Figure S4**).

## Discussion

This research develops novel understanding of the biomass and secondary production rates of fishes in reservoirs, with implications for the management of freshwater resources globally. Our estimates suggest reservoirs contain substantial pools of fish biomass comparable to other important values presented in the literature (**Table 2**) (34–37). Fish are core to food security and cultures in many nations across the world (1, 8, 38–40). While the literature has focused predominantly on the role of marine fisheries in food security, there is a growing recognition that inland fisheries play major and underappreciated roles (2, 17, 37, 41). In addition, we find non-linear trends in biomass exist both spatially and temporally. This suggests nuance exists in how reservoir production changes with time, and that reservoirs do not necessarily always experience steep declines in productivity over long time periods. Future work covering longer time periods is needed to better understand the potential for production declines with reservoir age.

**Table 2.**
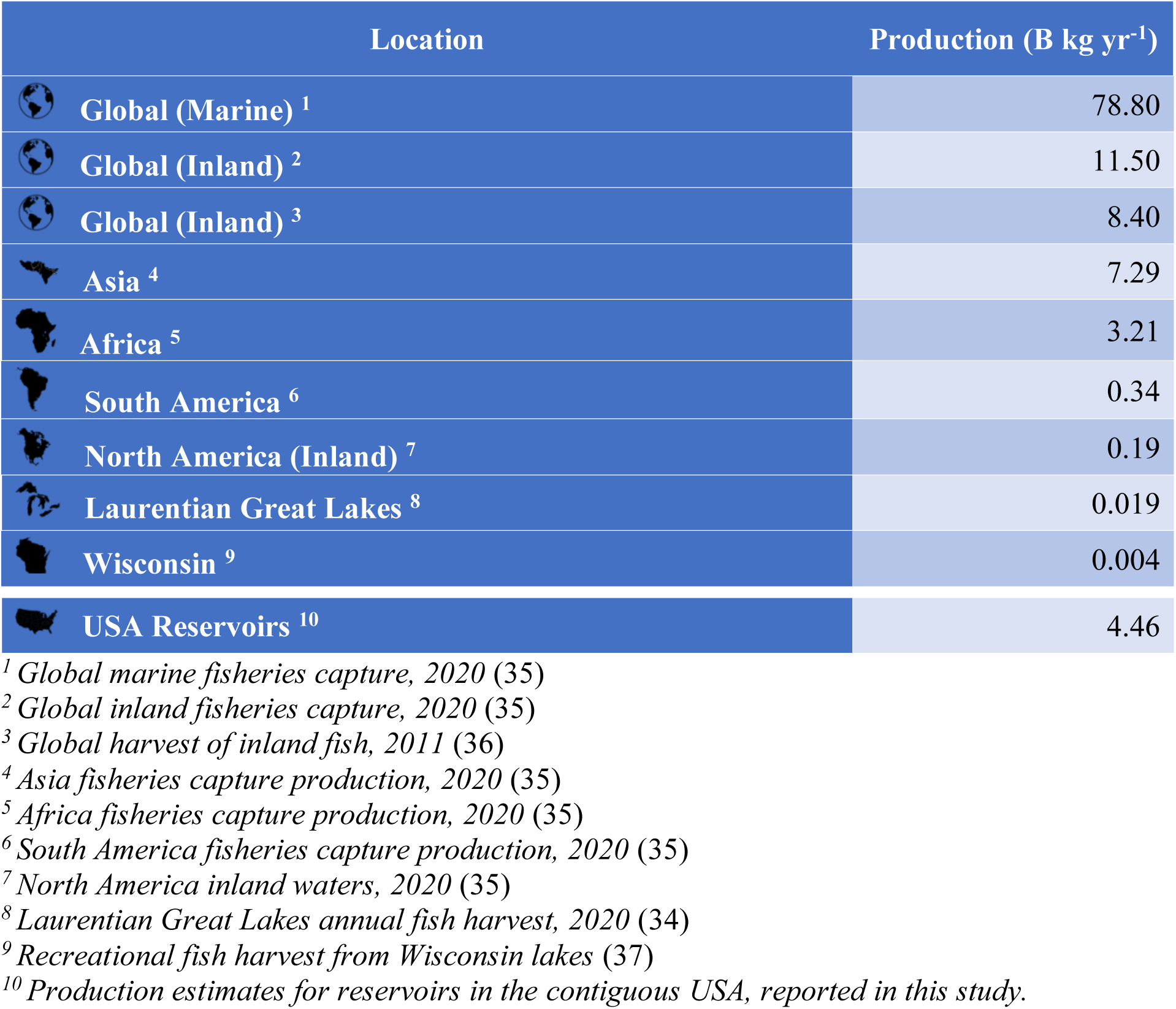
Current estimates of fish harvest and capture production from the literature compared with the results of this study.

| Location | Production (B kg yr<sup>-1</sup>) |
| --- | --- |
| Global (Marine) <sup>1</sup> | 78.80 |
| Global (Inland) <sup>2</sup> | 11.50 |
| Global (Inland) <sup>3</sup> | 8.40 |
| Asia <sup>4</sup> | 7.29 |
| Africa <sup>5</sup> | 3.21 |
| South America <sup>6</sup> | 0.34 |
| North America (Inland) <sup>7</sup> | 0.19 |
| Laurentian Great Lakes <sup>8</sup> | 0.019 |
| Wisconsin <sup>9</sup> | 0.004 |
| USA Reservoirs <sup>10</sup> | 4.46 |

<sup>1</sup> Global marine fisheries capture, 2020 (35)
<sup>2</sup> Global inland fisheries capture, 2020 (35)
<sup>3</sup> Global harvest of inland fish, 2011 (36)
<sup>4</sup> Asia fisheries capture production, 2020 (35)
<sup>5</sup> Africa fisheries capture production, 2020 (35)
<sup>6</sup> South America fisheries capture production, 2020 (35)
<sup>7</sup> North America inland waters, 2020 (35)
<sup>8</sup> Laurentian Great Lakes annual fish harvest, 2020 (34)
<sup>9</sup> Recreational fish harvest from Wisconsin lakes (37)
<sup>10</sup> Production estimates for reservoirs in the contiguous USA, reported in this study.

Our final reservoir classification schema is likely useful for future conservation science. Fish biomass and production data comport well with our classification schema, demonstrating that the ecology of reservoirs varies strongly alongside the reservoir classes. While in this study the classification schema was used to understand patterns in fisheries and food security, our classification may have additional applications towards effective reservoir management for the good of humanity, biodiversity, and fisheries conservation. For example, these classifications could be used in the study and management of limnology, food web ecology, and ecosystem dynamics of reservoirs throughout the USA. A particular advantage is that this classification system can be used at the national (USA) and regional (state or lower) scales, and thus may be of interest to a diversity of managers and scientists. Further, the same framework could also be applied globally or in any region where reservoir discharge and volume data are available.

The implications of large pools of fish biomass in reservoirs are severalfold. **Firstly,** an abundance of fixed carbon in resident reservoir organisms suggests a major and increasing role for reservoirs in the global carbon and freshwater cycles. Understanding the scope of this effect should attract research attention going forward. **Secondly,** it is clear from the magnitude of our estimates, that reservoirs (and probably other novel ecosystems) harbor additional sources of fish protein that are likely already being utilized substantively by societies. Awareness around this topic is highly limited within the ecological and social sciences. Yet without proper management, freshwater populations can quickly deteriorate and even collapse (1, 42). Therefore, one implication of our findings is that reservoirs globally would benefit from increased management attention. Without proper management, ecosystem services will be extremely limited, or in worst cases, just collapse. **Finally,** we note that reservoirs can also represent important habitats for native species, e.g., those resilient to fragmentation by dams (25, 43), or as potential novel habitat for species vulnerable to collapse in their native range (44, 45). There may be opportunity to craft reservoir ecosystems into emergency rooms for a subset of native species. However, conservation management of reservoirs to this point has generally not embraced this potential.

Reservoirs occupy a massive geographic footprint on the planet; thus, pools of fisheries biomass within reservoir ecosystems are relevant at all scales. While impoundment of rivers can create short-lived production spikes, these effects notoriously dissipate (46, 47), and long-term declines in production is a growing concern (27, 28). Similarly, global inland fisheries catch has the potential to either increase or decrease in response to climate change impacts, largely owing to variation in land-use surrounding individual waterbodies (2). Yet we observe that these production trends can have substantial spatiotemporal variation. Therefore our method of exploring and quantifying macroecological production patterns can aid in illuminating shifts in productive capacity, which in turn can be useful for conservation practitioners. Further, climate change has the potential to expedite or slow the rate of functional aging in existing reservoirs, and this topic is of growing interest (29). We do not suggest that reservoirs boost ecological or economic production; indeed construction of new dams will have net negative effects on ecosystems and surrounding communities (48–50). For example, available data suggest biomass and fish biomass and production rates were substantially higher in unregulated rivers prior to dams compared to reservoirs following impoundment (51).

Owing to their massive footprint, reservoir ecosystems now support globally relevant pools of fish biomass. Understanding the distribution and dynamics of this pool may be highly relevant from a global carbon standpoint. We show fish in USA reservoirs alone contribute 0.0045 Pg yr^-1^ of production (i.e., 4.5 kg yr^-1^). Reservoirs and lakes comprise 0.28 Pg C y^−1^ and 0.11 Pg C y^−1^ of the global carbon cycle, respectively (52). Although this quantity may appear small, it is on the scale so as to be relevant from a carbon cycle perspective. Furthermore, fish consumers classically exert control over food webs via trophic cascades, such that even a relatively small number of fish, or change in numbers, can play disproportionately impactful roles in carbon flux, nutrient cycling, and energy transfer (53–55). Indeed our biomass and production estimates may also represent partial indicators of ecological resilience (56, 57), especially when used in conjunction with local-scale functional diversity and food web metrics (58–60). Therefore, we encourage future freshwater scientists to make use of the reservoir classification framework and supplementary datasets (**Dataset S3**) developed in this study for other endeavors.

We note our method for estimating fish biomass is only one approach to generating such estimates, and we view these estimates as a valuable preliminary framework. For example, Deines et al. (36) utilized remotely-sensed chlorophyll concentrations from 80,012 lakes to approximate global lake fish harvest. In their approach, chlorophyll data were related to empirical estimates of fisheries harvest on a subset of lakes and these relationships were used to extrapolate fish harvest at scale. Similar methods have been used to assess production of terrestrial plants and other aquatic organisms (61, 62). Yet this method involves a key assumption that food web pathways of carbon transfer in aquatic ecosystems are roughly fixed relative to the “greenness” of the water. Increasingly, we understand many aquatic food webs are benthic in their functionality (63), which presents issues for remotely-sensed models of fish production based solely on “greenness”. Furthermore, there is less need to model biomass and production if these same data can be empirically-derived and are available (64, 65). One issue with our method is the limited spatial extent of the biomass surveys – because fish poisoning surveys were so heavily concentrated in southern USA states. Future studies aimed at reconciling fish biomass and production estimates using a variety of methodologies could be valuable. Furthermore, additional research might explore the sensitivity of estimates to varying littoral/pelagic habitat ratios (66). Ultimately, our estimates of standing stock biomass are probably conservative. The NID database is likely missing millions of smaller impoundments that escape local and federal regulation and thus inclusion in the NID. Small reservoirs, like small lakes, are numerous and notoriously difficult to inventory (67, 68). If these values were added, standing stock totals would only increase. However, most of our empirical biomass data were derived from larger reservoir environments, and limnetic extent is one of the primary drivers of the total standing stock calculation. Therefore it is likely that total standing stock values from these many smaller reservoirs would ultimately be small, even in aggregate, and may not change the total estimate appreciably (66, 69).

Reservoirs are important ecosystems to study further and to sustainably manage at all scales. There is near-complete regulation of the world’s rivers by widespread dam installation (19, 22, 70). Ecological effects of dams have been overwhelmingly negative and represent one of the principal drivers of freshwater biodiversity loss at all scales (20, 71). In addition, only 37% of Earth’s rivers (>1,000 km) maintain their natural flow without dam obstruction (72). Paradoxically however, little research has occurred on the novel ecosystems and changes to production left in the wake of dams. In many locations, reservoirs and fragmented rivers are the only freshwater ecosystems remaining (24); thus improved understanding of the ecology of these environments and their fisheries should be of interest to conservation scientists at all scales. Even though reservoirs are human-dominated environments, their global geographic footprint is testimony to their modern scope of importance. Taking down dysfunctional dams combined with improved management of remaining dams and reservoirs may represent a path towards improved freshwater fisheries, conservation, and food security. We encourage conservation scientists around the globe to rethink the potential for reservoirs to meet human-and conservation-based goals.

## Methods

For a full description of the methods see SI methods.

### Fish Biomass

Empirical measurements of fish biomass are rare (73). For most of the last century, it was common practice to use toxicants for sampling fish populations and community biomass, particularly in reservoir environments of southern USA (47, 74). Rotenone – a plant extract, was the primary chemical used in fish poisoning surveys. Rotenone kills fish by blocking oxygen uptake; thus, suffocating fish. While lethal, it is widely recognized in the fisheries literature as being one of the best methods for obtaining empirical fish biomass values (75). In surveys, block nets are used to isolate coves or other pelagic areas, and the poison pumped at appropriate concentration to kill all fishes present within the water column. During the 1970s, the US Fish and Wildlife Service launched the National Reservoir Research Program (NRRP), which as part of its mission, began collating prior rotenone surveys collected by other state and federal agencies and coordinating future surveys in USA reservoirs. Until now, the data from these efforts have only been available on paper.

We digitized the legacy National Reservoir Research Program rotenone (poisoning) fish biomass dataset and make these data publicly available as part of this paper (**Dataset S1**). The biomass data used for this study were generated from once widespread rotenone sampling programs which are now mostly banned (42). For environmental and humane reasons, sampling with toxicants has become rare over time and was never widely used in countries outside the USA. Due to rotenone’s efficacy, these rotenone datasets likely represent the best available, and most accurate, data on fish biomass in reservoir ecosystems to date; data of this kind are unlikely to ever be collected again. In total, the digitized dataset contains fisheries biomass data from 1,127 rotenone surveys on 301 USA reservoirs, 1948 – 1978, and spans twenty-two states (AL, AR, FL, GA, IA, IL, IN, KS, KY, LA, MA, MD, MO, MS, NC, NM, OK, SC, TN, TX, VA, WV). Species-specific biomass data are available; however, these data are yet to be entered into this database.

We used previously published data to adjust biomass estimates to account for biases associated with ineffectiveness of block nets and incomplete recovery of fish (75). **Adjustments** involved calculating an average of species recovery values presented in Table 10.1 of Shelton and Davies 1983 (75), and multiplying all reservoir biomass estimates by this constant (1.773056). Empirical fish biomass values were **joined to the open-access Omernik ecoregion dataset** (76). At its coarsest, level I, North America is subdivided into 15 ecological regions, level II into 52 regions, and level III into 104 regions. We used Omernik level II resolution for the purpose of this analysis, however, use of any Omernik level resolution resulted in similar biomass predictions. Finally, our biomass data were joined to the **2018 National Inventory of Dams** (NID) (77) containing 91,468 rows of data on large, regulated dams and their reservoirs in the United States. The NID dataset is the most complete dataset on the inventory of dams and their reservoirs known in the USA, though there are numerous (hundreds-of-thousands to millions) of small dams and other structures which are not captured through the NID. NID reservoirs were also joined to Omernik level II ecoregions. During our analyses, we identified some issues with the NID dataset that required action. For example, some larger reservoirs have multiple dams; thus, data were cleaned using coding rules that, to the best of our ability, ensured each reservoir was only being counted once. Also, some of the largest waterbodies in the NID are natural lakes with small dams (e.g., Lake Superior) and needed to be removed prior to analysis. Finally, reservoirs without geographic coordinates, ecoregion assignments, and missing surface area information needed attention prior to analysis. The tidied NID dataset used in this analysis held 85,470 rows. See SI methods and supporting R code for details on data cleaning and preparation.

### Reservoir Classification System

We developed a series of reservoir classification systems of increasing complexity using reservoir volume, discharge, and Omernik ecoregions. In our most refined classification system, which may be of interest to future researchers of USA reservoirs, we used a hierarchical approach to classification whereby reservoirs were grouped by their membership in Omernik’s level II ecoregions. Then for each ecoregion, we ran a k-means cluster analysis using reservoir maximum discharge and storage volume (log(x+1) transformed and scaled). Parallel with Rypel et al. (78), our reservoir classification was *a priori* constrained to four clusters for each ecoregion (i.e., large-slow, large-fast, small-slow, small-fast). K-means data clustering is a technique that scales well to large datasets and offers the advantage of flexibility, guaranteed convergence, tight clusters, and better interpretability for later re-use. We also explored other statistical classification algorithms, but none seemed to greatly augment results; we present here the results of our more straightforward clustering (**Dataset S2**).

### Reservoir biomass and production estimates

We developed five different reservoir classification schemas and thus five separate biomass estimates, allowing for some estimation of uncertainty (**Figure S1)** (79). Most available empirical biomass data were collected in twenty-two southern USA states, therefore, we calculate summary statistics for the southern USA as a sample-rich region, but also present extrapolations for the contiguous USA, while recognizing there are regional differences in the dynamics of fish biomass production.

Generalized Additive Mixed Models (GAMMs) were first created to examine biomass as a function of reservoir age under each of the five classification methods (80). GAMMs were fit using restricted maximum likelihood (REML) smoothness selection, Gamma family, and log link function (81). The two continuous predictors used in the models, *Reservoir-Age* and *Year-Sampled,* received thin plate spline smooths, the reservoir (*Ecosystem*) name received a random effect smooth, and *Classification* received smooth factor interaction for each of its categorical variables to determine whether smoothed fits varied by subclass. Classes with fewer than five data points were removed prior to running the respective model. Model quality was assessed via model convergence, basis checks, residual and partial residual plots, model summaries, and using second-order Akaike information criterion (**Table S1**). While deviance explained by the model is viewed as a more appropriate goodness-of-fit indicator for non-normal errors in non-gaussian models (81), both percent deviance explained and adjusted R^2^ are presented in **Table S1**.

Each model was used to independently **predict** fisheries biomass data beyond the final year of empirical biomass data (1978–1993) to standardize for noise resulting from reservoirs having been sampled at different points in time, and to estimate potential change in fisheries biomass within reservoirs over time (**Figure S2)**. Finally, to assess the model’s predictive ability, trends in empirical and predicted fish biomass over time in study reservoirs were examined and validated, as suggested by Pedersen et al. (80) (**Figure S3-S4**; see SI for validation techniques). Thus, results from Schema 5’s nearest reliable year (1993) were used to create total standing stock and production estimates, and main manuscript figures.

In each **calculation** method, class-specific averages of fish biomass were assigned as fish biomass estimates for any reservoir of the same class that did not have empirical rotenone data (**Table S2**). When no biomass estimates were available for an entire class, we substituted mean biomass across all sampled reservoirs. Once all reservoirs had been assigned a biomass estimate, reservoir biomass (kg ha^-1^) values were multiplied by the surface area of the reservoir (ha), or approximated surface area if none previously existed, to obtain a total standing stock (kg) estimate for every reservoir in the NID (**Dataset S3**). We then summed total standing stocks across the entire cleaned NID dataset to estimate total standing stock in southern USA and USA reservoirs for that classification approach. Finally, we also summed total standing stock by US state to highlight general geographic patterns. Fish production rates were estimated based on published production to biomass (P/B) ratios for whole fish communities from the literature (see SI Methods).

We provide additional details on summary analyses and validation procedures in the supplementary text. All cleaning and analytical code used R software (R version 4.3.0, R Core Team 2023) and is freely available and presented as part of this paper (https://github.com/caparisek/res_biomass_USA; DOI 10.5281/zenodo.8316696). All data and reservoir classifications are available on Zenodo (DOI 10.5281/zenodo.8317007).

The following R packages were used for this analysis: {tidyverse} v2.0.0 (82), {sf} v1.0.12 (83, 84), {sp} v1.6.0 (85, 86), {ggspatial} v1.1.8 (87), {tigris} v2.0.3 (88), {mgcv} v1.8.42 (81, 89–92), {MuMIn} v1.47.5 (93), {lmerTest} v3.1.3 (94), {smatr} v3.4.8 (95), {fBasics} v4022.94 (96), {data.table} v1.14.8 (97), {scales} v1.2.1 (98), {patchwork} v1.1.2 (99), {cowplot} v1.1.1 (100), {LaCroixColoR} v0.1.0 (101).

## Acknowledgements

CAP was supported by the UC Davis Center for Watershed Sciences’ Bechtel Next Generation Funds. ALR was supported by the Agricultural Experiment Station of the University of California, Project CA-D-WFB-2467-H, and by the California Trout and Peter B. Moyle Endowment for Coldwater Fish Conservation. This material is also supported by the National Science Foundation (NSF) under Grant DEB-2225284. Data were originally collated in paper form by Dr. Robert Jenkins, National Reservoir Research Program, U.S. Fish and Wildlife Service, with the cooperation of numerous state and federal land management agencies across the U.S; we thank all these biologists for their major contributions and efforts.

## Supporting Information for

## Methods

### NID Preparation

Data were cleaned using rules to remove duplicate rows representing multiple dams on the same reservoirs in the National Inventory of Dams (NID) dataset (1). We identified the 100 largest reservoirs and manually removed those which were clearly natural water bodies (n = 25; e.g., Lake Superior (MI), Lake Winnebago (WI), Mille Lacs Lake (MN), Clear Lake (CA)), as these would significantly and artificially inflate total standing stock and secondary production (hereafter, “production”) estimates for this reservoirs study.

Reservoirs with no latitude or longitude information (n = 242) in the NID received coordinates for its county’s centroid; of these, reservoirs without county information (n = 16) were manually located using other NID characteristics and supplied with coordinates that associated them with their correct Omernik ecoregion. Likewise, reservoirs whose precise latitude-longitude coordinates fell outside an ecoregion polygon (e.g., was implausibly located in an ocean; n = 27), thus receiving an N/A ecoregion after spatial joining, were manually assigned their correct ecoregion.

Reservoirs without surface area information (n = 18,145) received an approximated surface area decided by the most common surface area for that Omernik level II ecoregion; this was done by taking the *mean(log(surface_area))* and back-transforming (*10 ^*). This approximated surface area column was only used in the final step of the total standing stock calculation when converting kg/ha to kg would have resulted in the loss of ∼21% of final results, otherwise, the original NID column was used as needed during analysis and reservoirs with missing surface area data received rules outlined in **Table S2**.

Finally, in two instances, empirically sampled reservoirs from the National Reservoir Research Programs (NRRP) had dams relocated or otherwise adjusted such that the same dam NID ID was associated with two unique reservoir ecosystems that were sampled at different points in time. The NID ID for these ecosystems received the suffix *A* or *B* in both the NRRP and NID to complete spatial joining prior to analysis.

### Predicted Fish Biomass

We performed several checks to determine an adequate breaking point in fish biomass predictions outside the sampling period such that they remain realistic, while also enabling standardization of reservoir biomass estimates to a common and more recent time period. **Estimated mean standard error (SE) of prediction fit** was calculated for each year. Predictions were capped at the year whose mean SE was no more than twice that of the last year empirical data had been collected. In this case mean SE in 1978 was 158.2, so predictions did not extend beyond 1993 (mean SE of fit 314.0). Likewise, **estimated mean standard deviation (SD) of prediction fit** yielded the same breaking point. Finally, a **broken-stick** (i.e., segmented) **regression** was performed as a final validation of this decision. Thus project trends are shown for 1978 – 1993 and projected biomass estimates from the nearest realistic year (1993) were used in calculations (**Figure S2**).

### Validation

We validated biomass predictions of all five models (1993) using an independent dataset, as recommended by Pedersen et al. (2). The validation dataset consisted of 42 independent reservoir fish rotenone survey biomass estimates (3, 4) that were not included in the legacy National Reservoir Research Program database. We matched validation ecosystems to our database and compared observed versus predicted total mass values using a mixed effect regression model where observed total fish mass in reservoirs was the dependent variable, predicted total fish mass (from our classification and biomass value assignment procedures) was the independent variable, and classification method was a random effect (**Figure S3**). We tested the slope of this relationship against a value of 1 using a t-test. We also compared the Schema 5’s predicted fish biomass for 1993 to NRRP empirical fish biomass to validate predictions (**Figure S4**).

### Fish production estimation

We estimated fish production based on biomass and fish community production to biomass (P/B) estimates from the literature and associated supplementary data. Secondary production rates in heterotrophic populations and communities are strongly predicted by biomass (5, 6). The statistical relationship between production and biomass is in fact a descriptor of the P/B ratio, which is mathematically the biomass turnover rate of the population or community (7, 8). Therefore, biomass can be multiplied by P/B to approximate production with a high degree of certainty (9).

Based on a meta-analysis of community fish production, biomass, and P/B values presented in Rypel and David 2017 (6), we estimate average community fish P/B for 149 aquatic ecosystems of similar latitudes to the USA is ∼1.3. Therefore, we multiplied our final biomass estimates by 1.3 to approximate annual production of fish communities in USA reservoirs.

**Figure S1.**
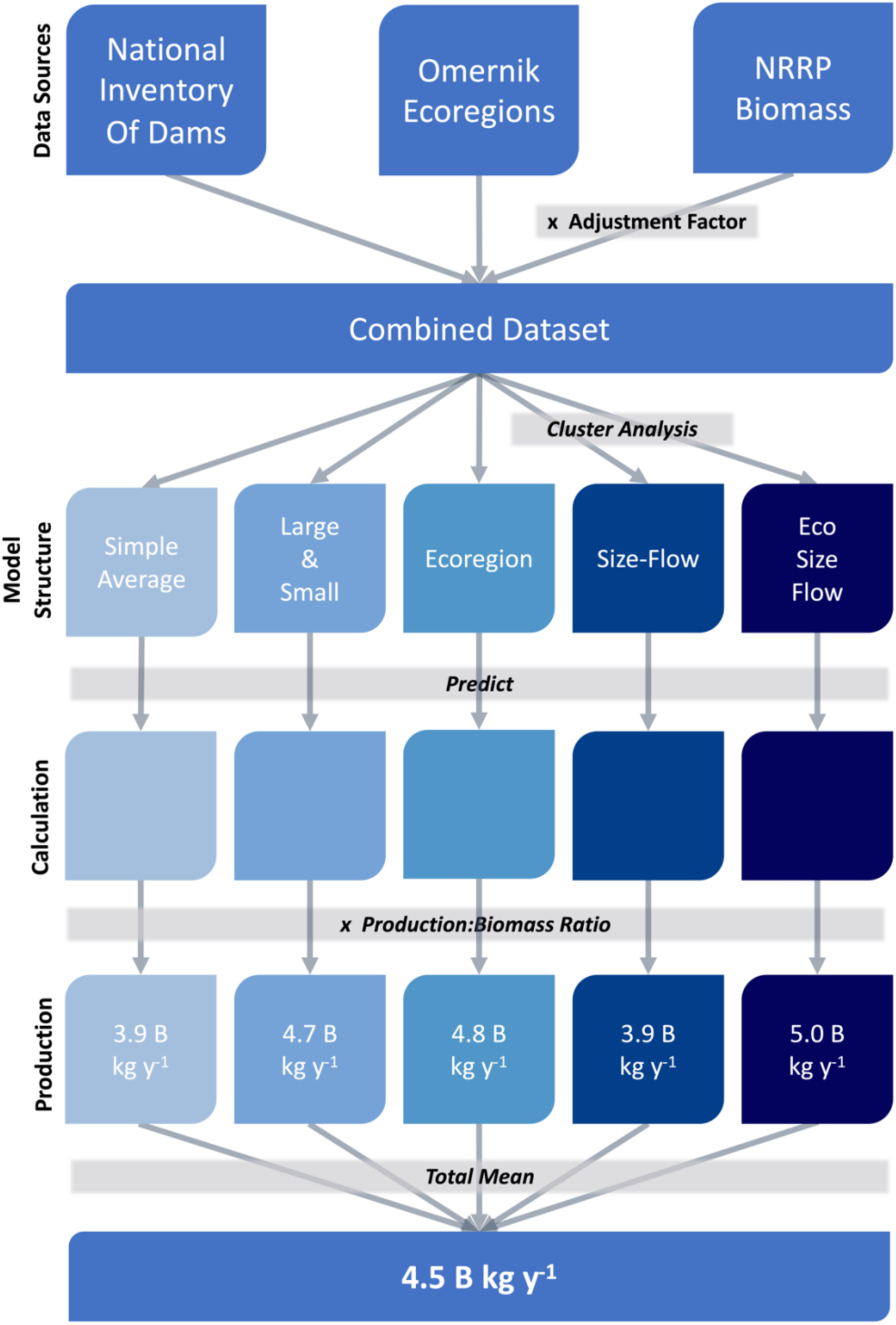
Workflow describing the development and processing of different data types in this study.

**Figure S2.**
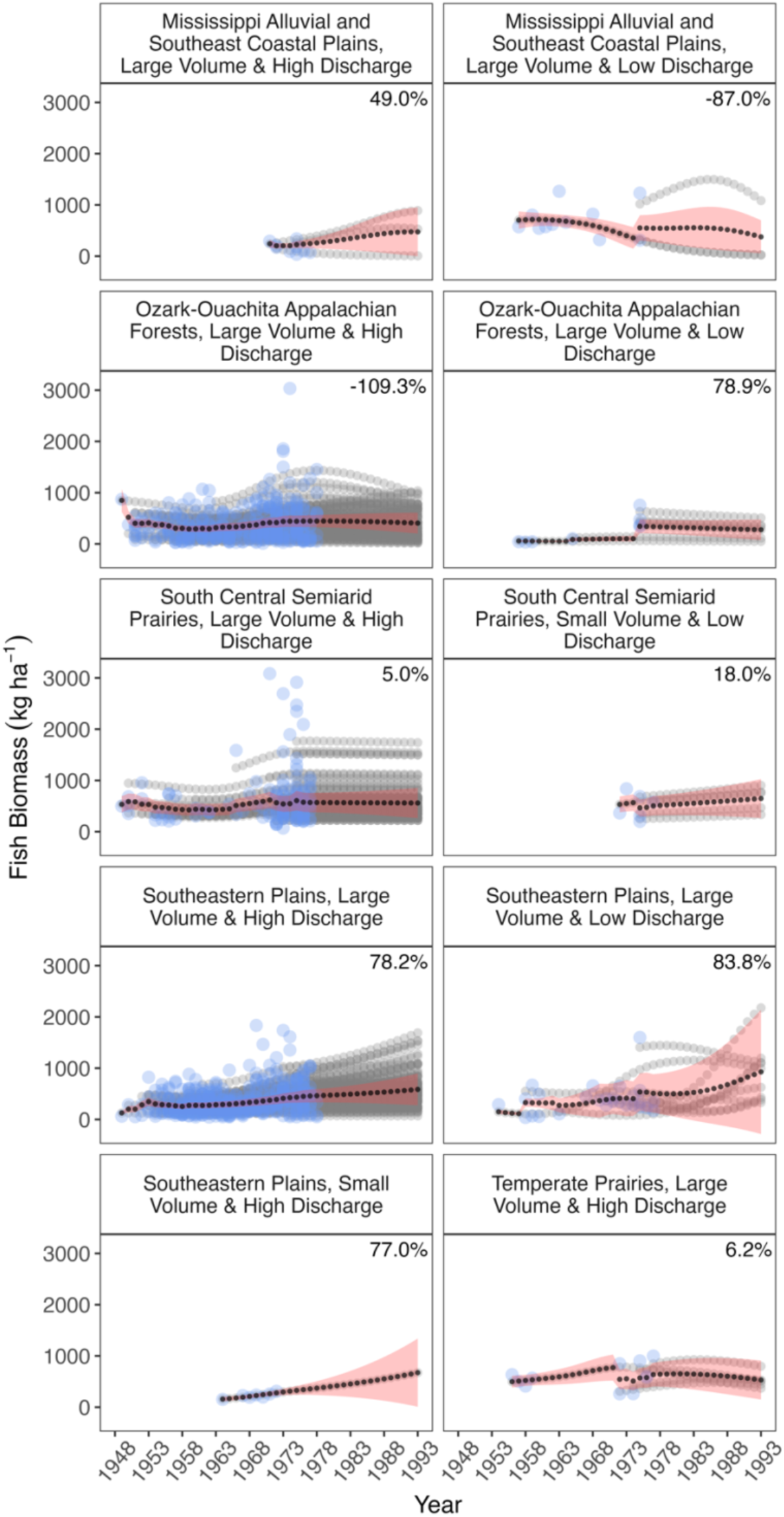
Temporal trends in empirical (blue) and predicted (gray) fish biomass over time in study reservoirs. Black line denotes mean of predictions from GAMMs using Schema 5 classification. Red ribbon denotes standard error of the model’s predicted fit. Percent change of fish biomass (kg/ha) from the initial year to the final (1993) is denoted in the upper right (((final year – initial year)/final year)*100).

**Figure S3.**
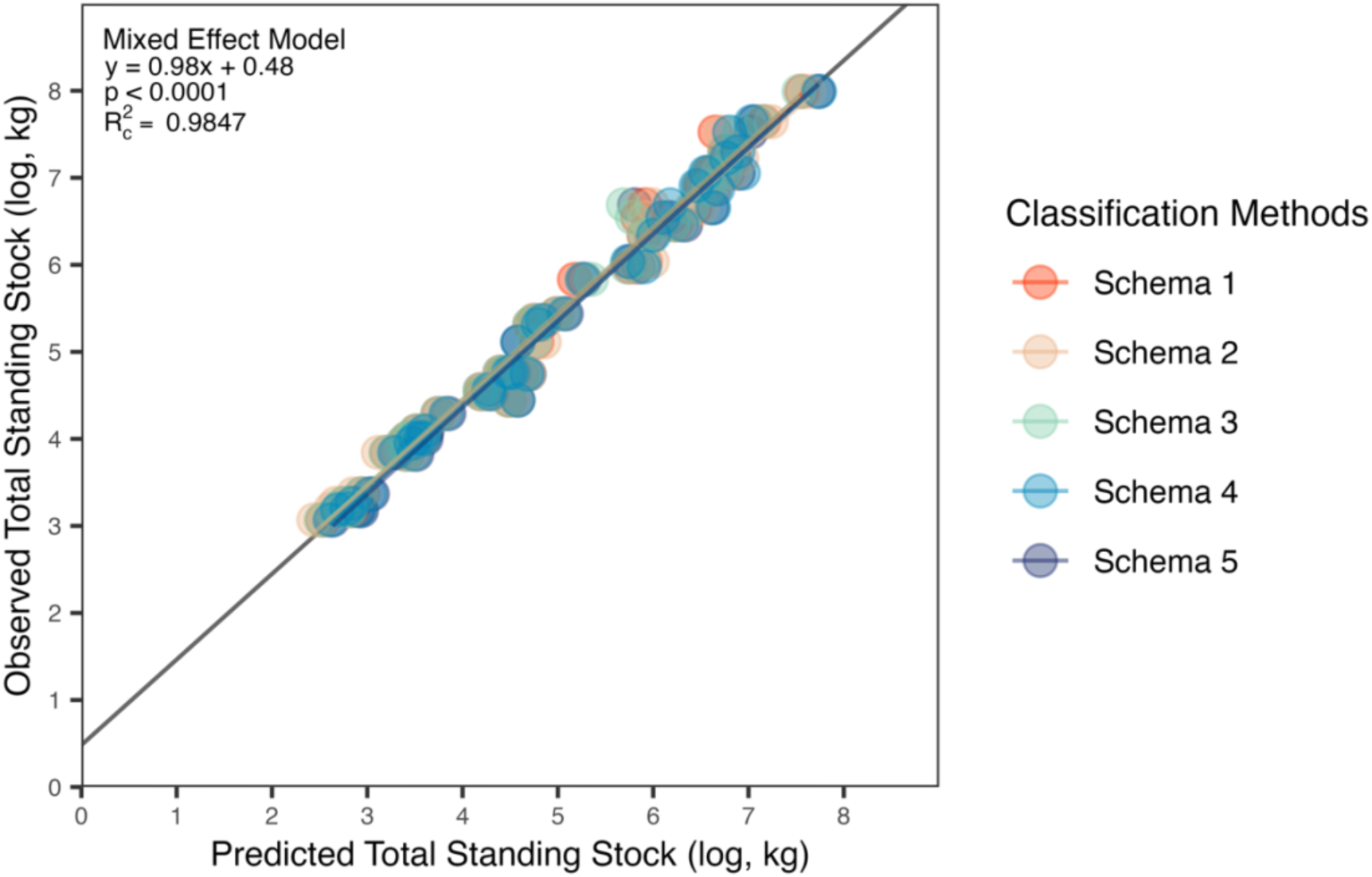
Validation analysis comparing predicted biomass values with an independent validation dataset from the literature on the same reservoirs. A mixed effect model was fit to the data with classification method as a random effect. Colored regression lines denote random effects and the thick dark line denotes the overall model regression.

**Figure S4.**
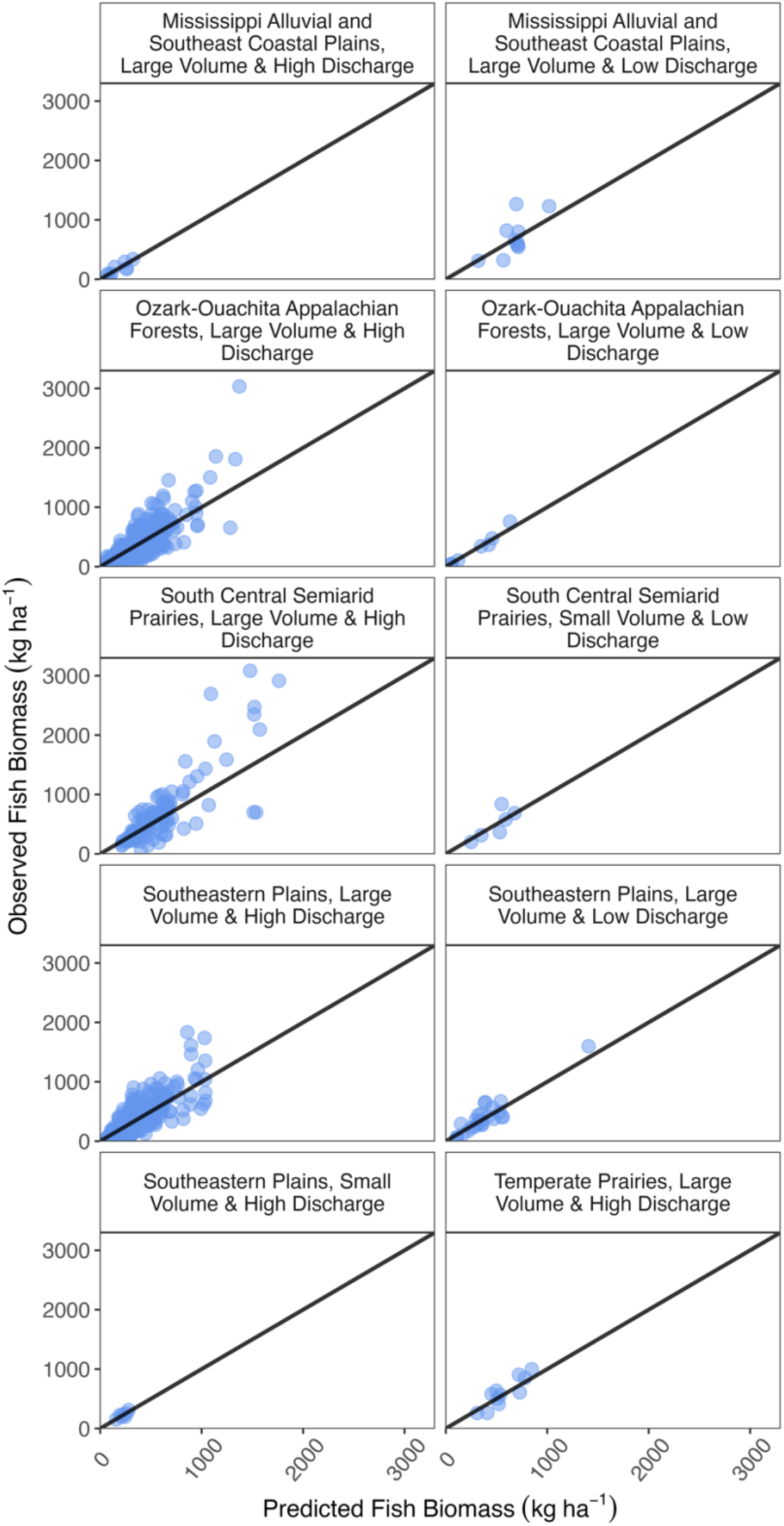
Observed fish biomass as a function of its predicted fish biomass. Black line illustrates a trend with slope = 1 and intercept = 0.

**Table S1.** GAMM model structure and summary output used for each of the classifications used for predictions. GAMMs used *Gamma(link = log)* and *“REML”* method. Smoothed variables are denoted within (*s*()). Smoothing is denoted with *bs =*.

| Schema | Model Structure | Deviance Explained (%) | Adjusted R-sq. | n | df | AIC |
| --- | --- | --- | --- | --- | --- | --- |
| – 1 –<br>Simple Average | Biomass ~<br>s(Reservoir_Age, bs="tp") +<br>s(Ecosystem, bs="re")+<br>s(Year, bs="tp") | 70.0 | 0.563 | 1,127 | 243.1628 | 14,646.54 |
| – 2 –<br>Large & Small | Biomass ~<br>s(Reservoir_Age, bs="tp",<br>by=Large_Small) +<br>s(Ecosystem, bs="re")+<br>s(Year, bs="tp") | 69.7 | 0.557 | 1,097 | 236.5298 | 14,274.33 |
| – 3 –<br>Size-Flow | Biomass ~<br>s(Reservoir_Age, bs="tp",<br>by=Cluster_Volume) +<br>s(Ecosystem, bs="re")+<br>s(Year, bs="tp") | 72.0 | 0.572 | 1,007 | 227.5092 | 13,052.05 |
| – 4 –<br>Ecoregion | Biomass ~<br>s(Reservoir_Age, bs="tp",<br>by=Ecoregion) +<br>s(Ecosystem, bs="re")+<br>s(Year, bs="tp") | 70.4 | 0.566 | 1,115 | 247.5736 | 14,485.05 |
| – 5 –<br>Eco-Size-Flow | Biomass ~<br>s(Reservoir_Age, bs="tp",<br>by=Eco_Size_Flow) +<br>s(Ecosystem, bs="re")+<br>s(Year, bs="tp") | 73.6 | 0.578 | 990 | 233.5400 | 12,786.79 |

**Table S2.** Five reservoir classification methods, ordered top-to-bottom from least complex to most complex.

| Schema | Classification methodology |
| --- | --- |
| – 1 –<br>Simple Average | A mean fish mass was calculated across all USA reservoirs. Reservoirs with no empirical biomass were given this mean. |
| – 2 –<br>Large & Small | Reservoirs were grouped into two classes, large and small. Median reservoir surface area for southern impoundments was 4.047 ha, and 4.856 ha for all USA impoundments; thus reservoirs below these thresholds were considered “small” while ones above were “large”. A mean fish mass was calculated for both large and small reservoirs and these values used for reservoirs where no biomass values were available.<sup>†</sup> |
| – 3 –<br>Size-Flow | Across all USA reservoirs, a cluster analysis constrained to four clusters was performed using reservoir volume (storage capacity) and maximum discharge. Mean fish mass was calculated across all reservoirs for a given cluster. Cluster mean fish mass values were used to assign a fish mass value to any reservoir missing empirical data. Clustering for the distinct southern and all USA reservoirs was performed separately.<sup>†</sup> |
| – 4 –<br>Ecoregion | Omernik level II ecoregions were identified for every reservoir. Mean fish mass was calculated across all reservoirs for an ecoregion. Ecoregion mean fish mass values were used for any reservoir with unknown fish biomass values.<sup>†</sup> |
| – 5 –<br>Eco-Size-Flow | Omernik level II ecoregions were identified for every reservoir. Within every ecoregion of the USA, a cluster analysis constrained to four clusters was performed using volume (i.e., storage capacity) and maximum discharge. Mean fish mass was calculated across all reservoirs for a given cluster. We used cluster mean fish mass values to assign a fish mass value to any reservoir missing empirical data.<sup>†</sup> |

<sup>†</sup>In the event there were no empirical data for a given cluster or ecoregion, the mean value across all USA reservoirs was used.

**Dataset S1 (separate file).** Digitized fish biomass data from National Reservoir Research Program rotenone project.

**Dataset S2 (separate file).** Reservoir classification summary statistics using NID maximum discharge (m^3^s), NID storage volume (m^3^), and Omernik level II ecoregions.

**Dataset S3 (separate file).** Reservoir classification and Schema 1-5’s predicted biomass (kg ha^-1^) and total standing stock (kg) linked to each reservoir in the refined version of the National Inventory of Dams database.

